# Wnt signaling recruits KIF2A to the spindle to ensure chromosome congression and alignment during mitosis

**DOI:** 10.1101/2020.12.22.404020

**Authors:** Anja Bufe, Ana García del Arco, Magdalena-Isabell Hennecke, Matthias Ostermaier, Anchel de Jaime-Soguero, Yu-Chih Lin, Anja Ciprianidis, Ulrike Engel, Petra Beli, Holger Bastians, Sergio P. Acebrón

## Abstract

Canonical Wnt signaling plays critical roles in development and tissue renewal by regulating β-catenin target genes. Recent evidence showed that β-catenin-independent Wnt signaling is also required for faithful execution of mitosis. This mitotic Wnt signaling functions through Wnt-dependent stabilization of proteins (Wnt/STOP), as well as through components of the LRP6 signalosome. However, the targets and specific functions of mitotic Wnt signaling still remain uncharacterized. Using phosphoproteomics, we identified that Wnt signaling regulates the microtubule depolymerase KIF2A during mitosis. We found that Dishevelled recruits KIF2A via its N-terminal and motor domains, which is further promoted upon LRP6 signalosome formation during mitosis. We show that Wnt signaling modulates KIF2A interaction with PLK1, which is critical for KIF2A localization at the spindle. Accordingly, Wnt signaling promotes chromosome congression and alignment by monitoring KIF2A protein levels at the spindle poles both in somatic cells and in pluripotent stem cells. Our findings highlight a novel function of Wnt signaling during cell division, which could have important implications for genome maintenance, notably in stem cells.

**SIGNIFICANCE:** Wnt signaling plays essential roles in embryonic patterning, stem cell renewal, and cell cycle progression from G1 to S phase via the regulation of *β*-catenin target genes. Here, we show that Wnt signaling also promotes faithful execution of mitosis by ensuring chromosome congression and alignment before cell division, including in pluripotent stem cells. We demonstrate that the Wnt signaling transducer Dishevelled recruits the mitotic kinesin KIF2A, and mediates its binding to the spindle. KIF2A is a microtubule depolymerase that controls chromosome alignment and congression during mitosis. Consequently, we found that inhibition of Wnt signaling leads to KIF2A-dependent chromosome congression and alignment defects.

## INTRODUCTION

The canonical Wnt signaling pathway plays essential roles in embryonic development and tissue homeostasis (Niehrs, 2010, Clevers et al., 2014). In particular, Wnt signaling governs stem cell maintenance and proliferation in many tissues, and its misregulation is a common cause of tumor initiation (Nusse and Clevers, 2017, Bugter et al., 2020).

Wnt ligands bind the Frizzled (FZD) receptors and the co-receptors low-density lipoprotein receptor-related proteins 5 and 6 (LRP5/6) (Tamai et al., 2000). The activated receptor complexes cluster on Dishevelled (DVL) platforms and are internalized via caveolin into endosomes termed LRP6 signalosomes, which triggers sequential phosphorylation of LRP6 by GSK3β and CK1γ (Davidson et al., 2005, Zeng et al., 2005, Yamamoto et al., 2006, Bilic et al., 2007, Cruciat et al., 2010). LRP6 signalosomes recruit the β-catenin destruction complex, which contains the scaffold proteins AXIN1 and adenomatous polyposis coli (APC), the kinases CK1α and GSK3β, and β-TrCP (Aberle et al., 1997). This recruitment inhibits the kinase GSK3β and releases β-TrCP, which leads to β-catenin stabilization and nuclear translocation in a IFT-A/KIF3A-dependent manner (Metcalfe et al., 2010, Taelman et al., 2010, Li et al., 2012, Kim et al., 2013, Vuong et al., 2018). LRP6 signalosomes mature into multivesicular bodies, sequestering the Wnt receptors together with GSK3β, thereby maintaining long-term activation of the Wnt pathway and promoting macropinocytosis (Taelman et al., 2010, Vinyoles et al., 2014, Kim et al., 2015, Garcia de Herreros and Dunach, 2019, Tejeda-Munoz et al., 2019, Albrecht et al., 2020). In contrast to Wnt ligands, the Wnt inhibitor DKK1 induces the clathrin-dependent internalization and turnover of LRP5/6 and thereby abrogates canonical Wnt signaling (Mao et al., 2001).

LRP6 signalosome formation peaks in mitosis (Davidson et al., 2009, Chen et al., 2018). On the one hand, the LRP6 competence to respond to Wnt ligands is promoted during G2/M by a priming phosphorylation at its intracellular domain by CDK14/16 and CCNY/CCNYL1 (Davidson et al., 2009, Niehrs and Acebron, 2012). On the other hand, CDK1 phosphorylates and recruits B-cell CLL/lymphoma 9 (BCL9) to the mitotic LRP6 signalosomes (Chen et al., 2018). BCL9 protects the signalosome from clathrin-dependent turnover, thereby sustaining basal Wnt activity on the onset of mitosis.

Mitotic Wnt signaling not only modulates β-catenin (Davidson et al., 2009), but piling evidence suggest that it promotes a complex post-translational program during mitosis (Acebron and Niehrs, 2016). For instance, we have shown that mitotic Wnt signaling promotes stabilization of proteins (Wnt/STOP), which is required for cell growth and ensures chromosome segregation in somatic and embryonic cells (Acebron et al., 2014, Huang et al., 2015, Stolz et al., 2015, Acebron and Niehrs, 2016, Chen et al., 2018, Madan et al., 2018, Hinze et al., 2019). Particularly, basal Wnt/STOP activity maintains proper microtubule plus end polymerization rates during mitosis, and its misregulation leads to whole chromosome missegregation (Stolz et al., 2015). Furthermore, mitotic Wnt signaling controls the orientation of the spindle (Kikuchi et al., 2010), and promotes asymmetric division in stem cells through components of the LRP6 signalosome (Habib et al., 2013). Accordingly, several Wnt components functionally associate with centrosomes, kinetochores and the spindle during mitosis (Fumoto et al., 2006, Hadjihannas et al., 2006, Kikuchi et al., 2010, Niehrs and Acebron, 2012). Consequently, both aberrant upregulation or downregulation of Wnt signaling have been associated with chromosome instability (CIN) (Fodde et al., 2001, Hadjihannas et al., 2006, Stolz et al., 2015), which is a hallmark of cancer (Kops et al., 2005). Despite the importance of these processes for tissue renewal and genome maintenance, the targets and specific functions of mitotic Wnt signaling remain largely uncharacterized.

Kinesin Family Member 2A (KIF2A) is a part of the kinesin-13 group (KIF2A,B,C) of minus end microtubule depolymerases (Manning et al., 2007, Walczak et al., 2013, Bakhoum et al., 2018). KIF2A is essential for the scaling of the spindle during early development (Wilbur and Heald, 2013), and plays critical roles in neurogenesis by modulating both cilium disassembly and neuronal wiring (Homma et al., 2003, Miyamoto et al., 2015, Ogawa and Hirokawa, 2015, Homma et al., 2018, Zhang et al., 2019). In dividing cells, KIF2A was originally thought to be required for the bipolar spindle assembly (Ganem and Compton, 2004), which was later proven to be due to an off-target effect (Tanenbaum et al., 2009). Current evidence supports a role of KIF2A in the regulation of microtubule dynamics at the spindle poles, thereby ensuring the congression, alignment and segregation of chromosomes (Uehara et al., 2013, Eagleson et al., 2015, Kwon et al., 2016, Yi et al., 2016, Ali et al., 2017, Xu et al., 2018). Genetic depletion of KIF2A in mouse leads to neonatal lethality and to severe brain malformations, including microcephaly (Homma et al., 2003, Poirier et al., 2013, Homma et al., 2018). KIF2A recruitment to microtubules is tightly coordinated by several protein kinases (Jang et al., 2009, Knowlton et al., 2009, Uehara et al., 2013, Miyamoto et al., 2015, Ogawa and Hirokawa, 2015, Ali et al., 2017, Trofimova et al., 2018, Xu et al., 2018). For instance, phosphorylation of KIF2A by Polo-like kinase 1 (PLK1) stimulates its recruitment to and activity at the spindle (Jang et al., 2009, Miyamoto et al., 2015, Ogawa and Hirokawa, 2015, Trofimova et al., 2018, Xu et al., 2018). On the other hand, Aurora kinase A and B inhibit KIF2A activity and restrict its subcellular localization during mitosis (Jang et al., 2009, Knowlton et al., 2009, Uehara et al., 2013).

Here, we show that mitotic Wnt signaling promotes chromosome congression and alignment in metaphase by recruiting KIF2A to the spindle in both somatic cells and pluripotent stem cells. We found that KIF2A is recruited by the LRP6 signalosome during mitosis. Mechanistically, we identified that KIF2A clusters with Dishevelled via the N-terminal and motor domains of the depolymerase. We show that Wnt signaling controls KIF2A interaction with PLK1, which is critical for KIF2A localization at the spindle poles. We propose that basal Wnt signaling ensures chromosome congression and alignment prior cell division by modulating the spindle minus-end depolymerization dynamics through KIF2A.

## RESULTS

### Mitotic Wnt signaling recruits KIF2A to the spindle

Mitosis is driven by post-translational modifications of proteins, most notably phosphorylation (Olsen et al., 2010). To identify direct targets and functions of Wnt signaling in mitosis, we performed a phosphoproteome-wide analysis using stable isotope labelling with amino acids in cell culture-based mass spectrometry (SILAC-MS) (Figure 1A). In detail, we synchronized SILAC-labelled immortalized RPE1-hTert (RPE1) cells at G1/S phase, released the cells into the cell cycle and harvested mitotic cells synchronized in mitosis with the mitotic kinesin-5 inhibitor Dimethylenastron (DME). Cells were treated for 1.5 h before harvesting with or without the canonical Wnt signaling inhibitor DKK1 (Figure 1A,B, S1A-C). Phosphorylated peptides were enriched from whole cell lysates, followed by their identification using ultra-high-performance liquid chromatography-tandem MS (LC-MS/MS).

**Figure 1.**
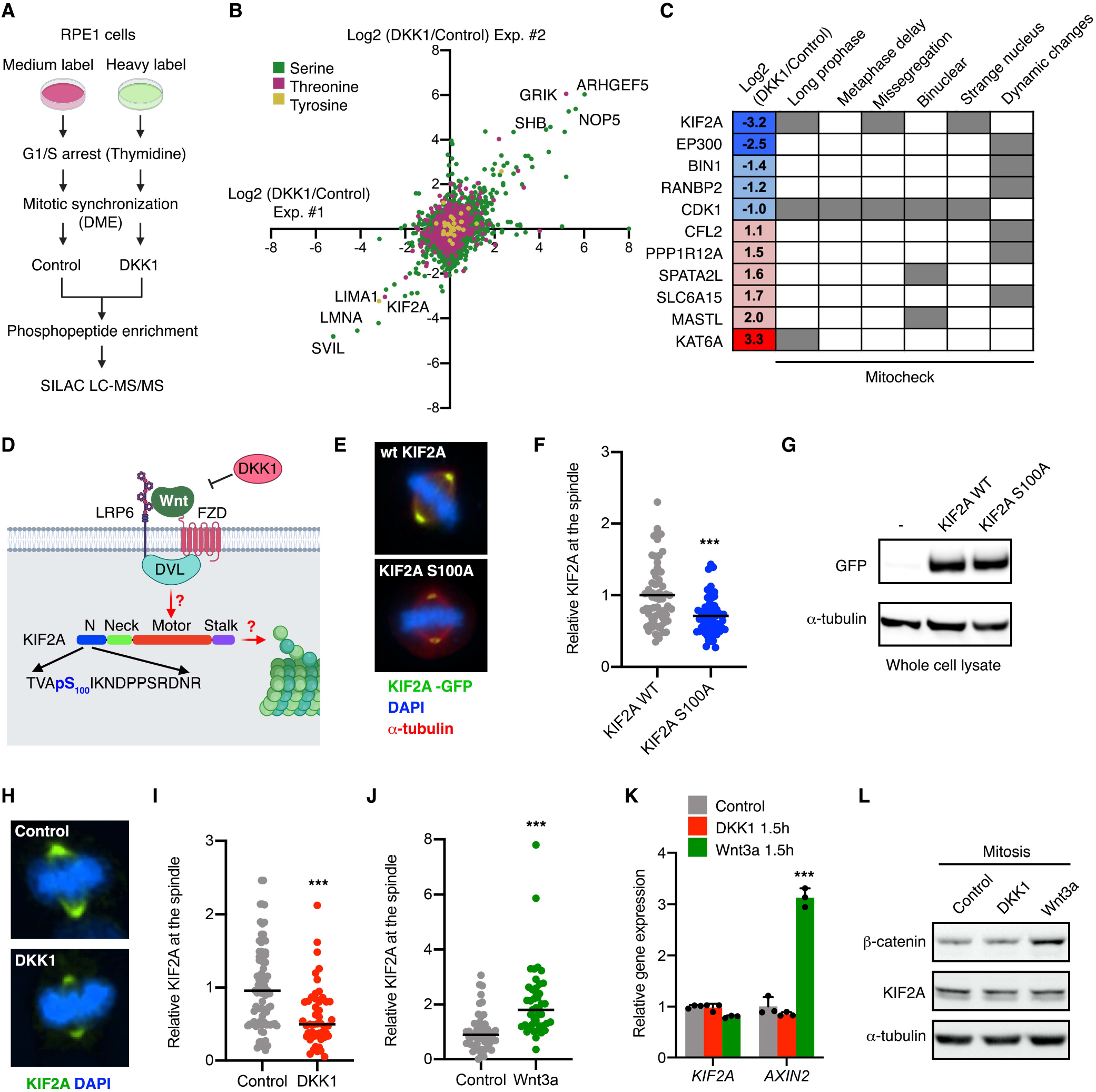
Mitotic Wnt signaling recruits KIF2A to the spindle. **A**, Scheme of SILAC MS analysis to identify changes in the phosphoproteome upon Wnt inhibition by DKK1 in mitotic cells. **B**, Scatter plot comparing the differential counts of phosphopeptides of two biological replicates. Axes show the log2-fold change in phosphopeptide counts between Control and DKK1 treatment in mitotic RPE1 cells. Phosphopeptides are color-coded for their modified residue. **C**, Table containing the log2-fold change in the phosphopeptide counts of the indicated proteins from our performed MS screen, and the mitotic phenotypes identified for their siRNAs in the Mitocheck database. **D**, Scheme showing the Wnt receptor complex and KIF2A. The phosphopeptide sequence identified in (**A**,**B**) is shown below with KIF2A phosho-S100, which is downregulated by DKK1, highlighted in blue. **E**, Representative immunofluorescence microscopy images showing overexpressed KIF2A-GFP protein levels (green) in HeLa cells during metaphase. **F**, Quantification of overexpressed KIF2A wt and S100A at the spindle of mitotic HeLa cells from (**E**). Mean of n > 61 cells per condition of a representative experiment of n = 3 independent experiments are shown. **G**, Representative western blots from n = 3 independent experiments showing lysates from HeLa transfected as indicated. **H**, Representative immunofluorescence microscopy images showing endogenous KIF2A protein levels (green) in RPE1 cells during metaphase treated as indicated 1.5 h before mitosis. **I**,**J**, Quantification of KIF2A at the spindle of mitotic RPE1 cells upon treatment for 1.5h with control or DKK1 conditioned media (**I**), or with control or Wnt3a conditioned media (**J**). In (**I**), mean of n > 43 cells per condition of a representative experiment of n = 3 independent experiments is shown. In (**J**), mean of n > 44 cells per condition from n = 2 independent experiments pooled is shown. **K**, qPCR analysis of *KIF2A* and *AXIN2* expression levels in G2/M arrested RPE1 cells upon 1.5 h treatment with the indicated conditioned media. Data is displayed as mean ± SD of 3 biological replicates. **L**, Representative western blots from n = 3 independent experiments showing cytoplasmic lysates from RPE1 cells synchronized in mitosis and treated for 1.5 h with the indicated conditioned media.

We quantified 12,208 phospho-sites in two independent replicate experiments (Figure 1B). We observed a high quantitative reproducibility between the replicate experiments (Figure 1B) and identified 21 upregulated and 73 downregulated phosphopeptides upon DKK1 treatment. Among the DKK1-regulated proteins, we found a moderate enrichment of proteins involved in cytoskeleton organization (p = 1.3⨯10^−4^) and regulation of cell cycle arrest (p = 6.7⨯10^−4^), suggesting an active role of Wnt signaling in mitotic progression (Figure S1D). To select Wnt target candidates with potential functions in mitosis, we examined the Mitocheck database (Neumann et al., 2010), which provides a comprehensive dataset of mitotic phenotypes obtained upon a genome-wide siRNA screen. Five of our candidates (KIF2A, KAT6A, MASTL, CDK1 and SPATA2L) displayed specific mitotic phenotypes (Figure 1C). For this study, we focused on KIF2A, which is required for chromosome congression, alignment and segregation of chromosomes (Uehara et al., 2013, Wilbur and Heald, 2013, Eagleson et al., 2015, Kwon et al., 2016, Yi et al., 2016, Ali et al., 2017, Xu et al., 2018).

Our mass spectrometry analysis revealed that DKK1 treatment reduced the phosphorylation of KIF2A serine 100 by ∼9-fold (Figure 1B-D). This uncharacterized phospho-site has been shown in previous proteomics analyses to be reduced upon PLK1 inhibition (Santamaria et al., 2011), and it is located in the N-terminal region, which modulates KIF2A recruitment to the spindle (Santamaria et al., 2011, Uehara et al., 2013, Ali et al., 2017) (Figure 1D). Notably, we found that mutation of serine 100 to alanine reduced KIF2A recruitment to the spindle, without affecting its total protein levels (Figure 1E-G). Hence, we examined by immunofluorescence whether Wnt signaling controls endogenous KIF2A localization during mitosis (Figure 1D). Importantly, inhibition of mitotic Wnt signaling in RPE1 cells by acute DKK1 treatment markedly reduced spindle-bound KIF2A during metaphase (Figure 1H,I), thereby phenocopying KIF2A S100A. Conversely, stimulation with Wnt3a during G2/M boosted the localization of endogenous KIF2A at the spindle (Figure 1J).

Mitotic Wnt signaling has been shown to regulate its targets by different mechanisms including via β-catenin-dependent transcription, Wnt-dependent stabilization of proteins (Wnt/STOP), and direct interactions with components of the LRP6 signalosome (Niehrs and Acebron, 2012, Acebron and Niehrs, 2016). Neither DKK1 nor Wnt3a treatment impacted KIF2A transcriptional or protein levels (Figure 1K,L). Furthermore, we have previously reported that Wnt/STOP regulates microtubule plus end dynamics in mitosis (Stolz et al., 2015). However, knockdown of KIF2A did not impact this function (Figure S1E). Taken together, these results exclude a role of Wnt/STOP and β-catenin in KIF2A regulation, and suggest that Wnt signaling directly promotes KIF2A phosphorylation and recruitment to the mitotic spindle.

### KIF2A is recruited by LRP6 signalosomes in mitosis

We next examined whether any Wnt signaling transduction components interact with KIF2A using proximity ligation assays (PLA) in HeLa cells (Figure 2A). We identified *in situ* KIF2A complexes with the signalosome scaffolding proteins AXIN1 at the spindle, and specially with DVL2 in the cytoplasm of mitotic cells (Figure 2B,C, complexes are shown in red). On the other hand, LRP6, GSK3β, CK1ε, β-TRCP and β-catenin only displayed background signal (Figure 2B,C). Importantly, co-expression of Wnt3a, FZD8, AXIN1 and LRP6, which induce LRP6 signalosomes (Figure 2D) (Bilic et al., 2007), boosted KIF2A recruitment to DVL2 during mitosis (Figure 2E,F). Furthermore, additional PLAs revealed *in situ* interaction between endogenous Dishevelled (DVL1-3) and KIF2A in the cytoplasm of mitotic cells, which was significantly impaired upon inhibition of mitotic Wnt signaling by DKK1 (Figure 2G,H).

**Figure 2.**
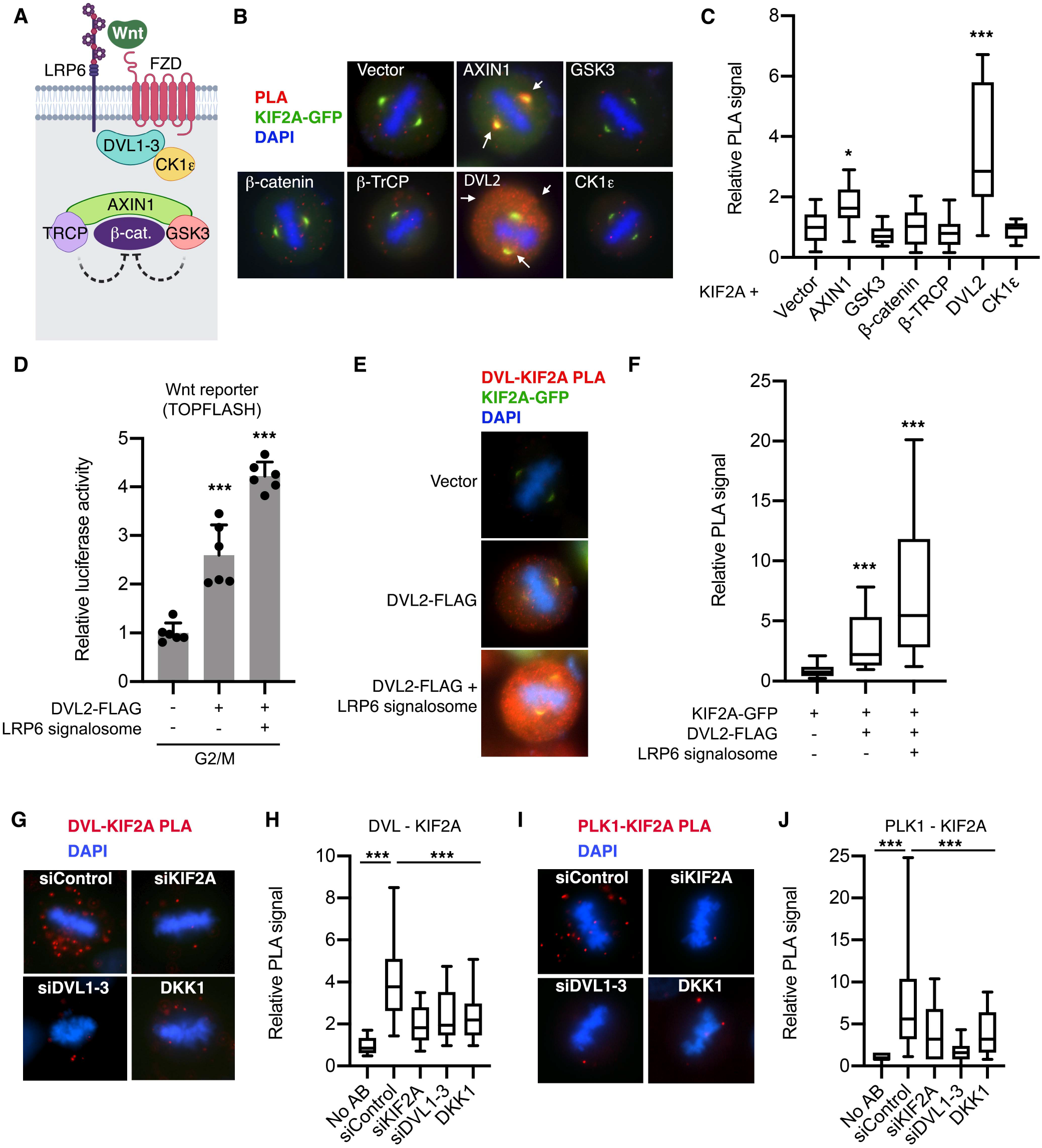
KIF2A is recruited by the mitotic LRP6 signalosome. **A**, Scheme of the Wnt signaling transduction components analyzed in this figure. **B**,**E**, Representative microscopy images of *in situ* Proximity ligation assays (PLAs) in mitotic HeLa cells transfected with GFP-KIF2A and the indicated constructs from n ≥ 2 independent experiments. LRP6 signalosome: Wnt3a/LRP6/FZD8/AXIN1. **C, F**, Box plot showing the quantification of the PLA signal from (**B**,**E**) and normalized to the empty vector control. Data is displayed as median PLA signal flanked by upper and lower quartile, followed by the 10-90% extremes of n > 18 mitotic cells in (**C**), or n > 65 mitotic cells in (**F**) from n ≥ 2 independent experiments. **D**, TOPflash reporter assays in HeLa cells upon co-transfection with empty vector, DVL2 or DVL2 together with the rest of the components of the LRP6 signalosome (Wnt3a/LRP6/FZD8/AXIN1). Data is displayed as mean ± SD of 6 biological replicates. **G, I**, Representative microscopy images of *in situ* Proximity ligation assays (PLAs) of endogenous proteins in mitotic HeLa cells. **H**,**J**, Box plot showing the quantification of the endogenous PLA signal from (**G**,**I**). Data is displayed as median PLA signal flanked by upper and lower quartile, followed by the 10-90% extremes of n > 26 mitotic cells representing n = 2 independent experiments.

To further validate the association between Dishevelled and KIF2A, we performed immuno-fluorescence experiments. Ectopic expression of Dishevelled leads to the formation of high molecular complexes through the DIX domains during interphase, which mimic LRP6 signalosomes (Bilic et al., 2007, Schwarz-Romond et al., 2007) (Figure S2A). Notably, DVL1-3 recruited KIF2A to their punctae (Figure S2A), further supporting a role of the signalosomes recruiting KIF2A through Dishevelled (Figure 2F). Next, we examined which domains of KIF2A are involved in its recruitment to the signalosomes (Figure S2B). Deletion of the KIF2A Neck or Stalk domains, which modulate the motor activity and dimerization, respectively (Trofimova et al., 2018), did not impair KIF2A localization in DVL2 punctae. On the other hand, deletion of the N-terminal (KIF2AΔN) or the motor domain (KIF2AΔmotor and KIF2A N-term) abolished the co-localization of KIF2A in DVL2 punctae (Figure S2C,D). Interestingly, both domains integrate signals that modulate KIF2A activity and localization, including by PLK1 (Jang et al., 2009, Santamaria et al., 2011, Uehara et al., 2013, Miyamoto et al., 2015, Ogawa and Hirokawa, 2015, Ali et al., 2017). Strikingly, S100A mutation largely abolished KIF2A recruitment to DVL2 punctae (Figure S2E,F), phenocopying the deletion of the whole N-terminal domain of KIF2A (Figure S2C). Taken together, we conclude that KIF2A requires its motor domain and the serine 100 at the N-terminal domain to functionally interact with Dishevelled.

Dishevelled is known to associate with PLK1 during mitosis (Kikuchi et al., 2010). Importantly, PLK1 inhibition reduces both KIF2A activity and S100 phosphorylation (Jang et al., 2009, Santamaria et al., 2011, Miyamoto et al., 2015, Ogawa and Hirokawa, 2015, Trofimova et al., 2018, Xu et al., 2018). Hence, we investigated whether Wnt mediates KIF2A interaction with PLK1. Notably, we found that knockdown of DVL1-3 or DKK1 treatment strongly reduced endogenous KIF2A association with PLK1 at the cytoplasm of mitotic cells (Figure 2I,J). Taken together, these results suggest mitotic LRP6 signalosomes recruit KIF2A and promote its interaction with PLK1.

### Wnt signaling promotes chromosome congression and alignment through KIF2A

Both the N-terminal and motor domains of KIF2A are required for its localization and activity (Uehara et al., 2013, Ali et al., 2017, Trofimova et al., 2018), and are essential for the functional interaction with Dishevelled (Figure S2C,D). Besides, our data indicate that loss of mitotic Wnt signaling reduces KIF2A association with PLK1 and to the spindle in metaphase (Figure 1F,G and 2I,J). Hence, we next investigated whether mitotic Wnt signaling controls chromosome congression and alignment through KIF2A.

Knockdown of the signalosome components LRP6 and Dishevelled (DVL1-3) impaired the recruitment of KIF2A to the spindle during metaphase in RPE1 cells (Figure 3A,B and S3A) similarly as we showed for DKK1 (Figure 1H,I). In agreement with the requirement of KIF2A in chromosome congression (Uehara et al., 2013, Wilbur and Heald, 2013, Eagleson et al., 2015, Kwon et al., 2016, Yi et al., 2016, Ali et al., 2017, Xu et al., 2018), inhibition of Wnt signaling by knockdown of LRP6 or Dishevelled, as well as by acute DKK1 treatment, led to congression and alignment defects as indicated by a wider chromosome spread parallel to the spindle poles (Figure 3A,C and S3B).

**Figure 3.**
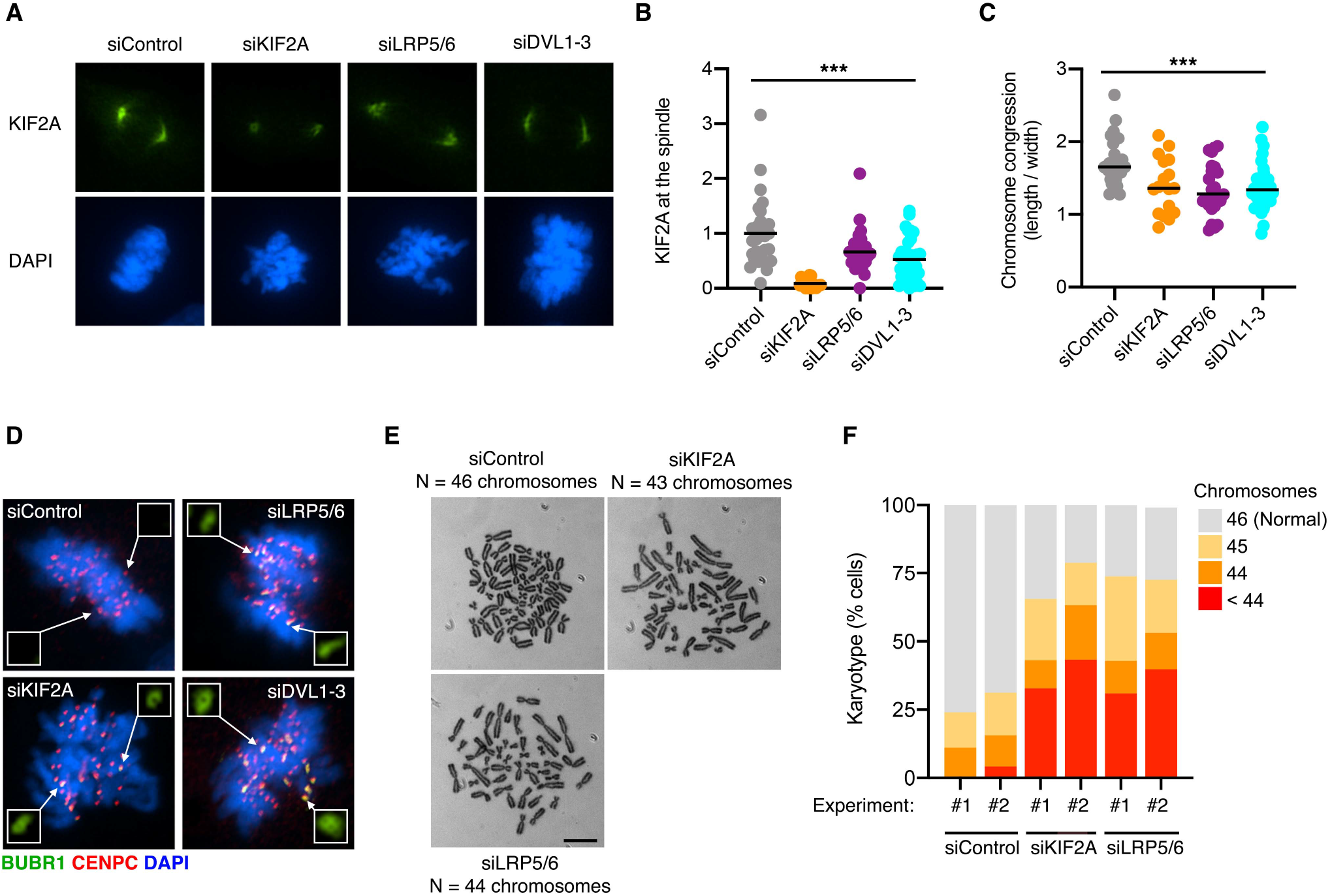
Basal Wnt signaling is required for chromosome congression and genome maintenance. **A**,**D**, Representative immunofluorescence microscopy images from n = 3 independent experiments showing mitotic RPE1 cells transfected with the indicated siRNAs and stained with antibodies against the indicated endogenous proteins. In (**D**), CENPC is used to mark the kinetochores and BUBR1 to highlight activation of the spindle assembly checkpoint (SAC) by misaligned chromosomes. **B**,**C**, Quantification of KIF2A immunofluorescence (Mean) and chromosome spread (Median) from (**A**). In (**C**), the length occupied by the chromosomes at the metaphase plate was divided by the width towards the spindle poles in n > 26 mitotic cells per condition. **E**,**F**, Chromosome number variability/aneuploidy of different cell clones derived from RPE1 cells transfected every 5 days with the indicated siRNAs and grown for 30 generations. In (**E**), representative examples of metaphase chromosome spreads, including normal (46, siControl) and aneuploid karyotypes (siKIF2A, siLRP6), are displayed. The graph in (**F**) shows the proportion of cells harboring a karyotype with chromosome numbers deviating from the modal in two experiments with 50 metaphase spreads per condition.

Failure to congress and align chromosomes can lead to chromosome instability (CIN) (Maiato et al., 2017). However, recent evidence on loss of KIF18A shows that chromosome misalignment does not necessarily lead to aneuploidy, unless kinetochore attachments are affected (Fonseca et al., 2019). We found that knockdown of KIF2A, or the signalosome components LRP6 and Dishevelled, resulted in recruitment of the spindle checkpoint protein BUBR1 to the kinetochores in metaphase (Figure 3D), which suggests a previous loss of tension at the sister kinetochores (Musacchio and Hardwick, 2002). Furthermore, using chromosome counting from metaphase spreads, we found that most cells (∼70%) lost one or more chromosomes upon prolonged loss of KIF2A or LRP6 over 30 days (Figure 3E,F). Hence, we conclude that both Wnt signaling and KIF2A are required for euploidy.

To further validate a role of Wnt signaling in promoting chromosome congression and alignment, we analyzed HeLa cells stably transfected with H2B-mCherry and EGFP-α-tubulin by live-cell imaging. Knockdown of KIF2A in HeLa cells led to severe chromosome congression or alignment defects in 24% of the divisions, compared with the 6% of the cases under control conditions (siControl) (Figure 4A-C, S3C). Inhibition of Wnt signaling in HeLa cells by siLRP5/6 or siDVL1-3 also resulted in chromosome congression or alignment defects in 25 and 30% of mitoses, respectively (Figure 4A-C, S3C). In addition, knockdown of KIF2A, LRP6 and DVL1-3 in HeLa cells delayed mitotic progression (Figure 4A,D).

**Figure 4.**
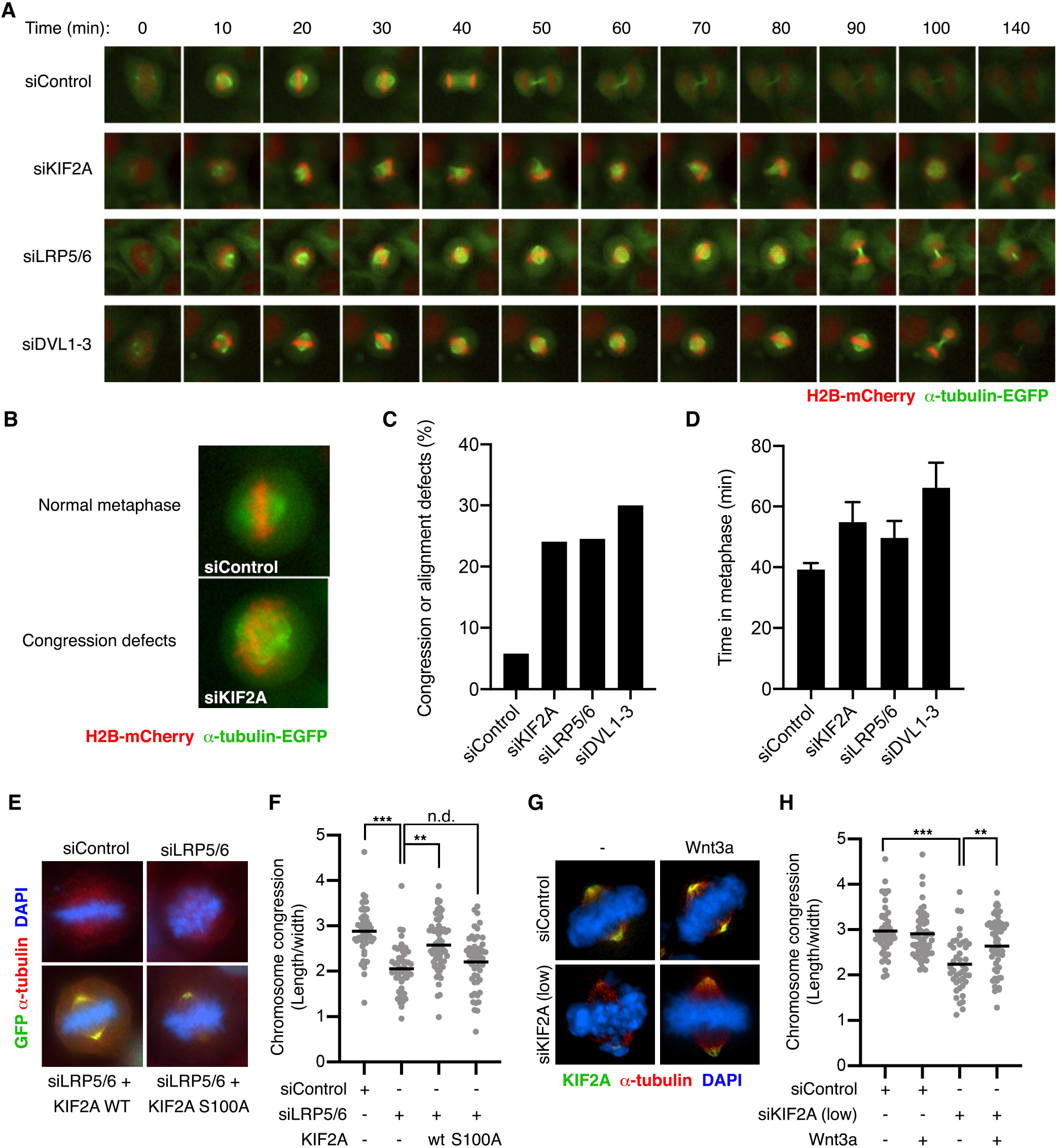
Wnt signaling promotes chromosome congression through KIF2A. **A**, Representative time lapses of mitotic HeLa cells stably expressing H2B-mCherry and EGFP-α-tubulin and transfected with the indicated siRNAs. **B**, Representative examples of metaphases with congressed chromosomes (Normal metaphases) and congression defects from (**A**). **C**, Live cell imaging analysis of mitoses displaying severe congression or alignment defects in HeLa cells from (**A**). Representative analysis of n > 50 cells per condition from one of n = 3 independent experiments. **D**, Analysis of mitotic progression in HeLa cells from (**A**). The average time from chromosome condensation to anaphase for all analyzed mitoses per condition is indicated in minutes. Data is displayed as mean ± SEM of n > 50 cells per condition from a representative analysis of n = 3 independent experiments. **E**,**G**, Representative immunofluorescence microscopy images of mitotic HeLa cells transfected with the indicated siRNAs and co-transfected or treated as indicated. **F**,**H**, Quantification of the chromosome congression from (**E**,**G**). The length occupied by the chromosomes at the metaphase plate was divided by the width towards the spindle poles in n > 35 mitotic cells per condition, and the median value is indicated. Representative images and analyses of n*≥* 3 independent experiments are shown.

Analyses of metaphases in fixed HeLa cells confirmed that inhibition of Wnt signaling by DKK1 or knockdown of LRP6 leads to congression and alignment defects (Figure 4E,F and S3D). Importantly, ectopic expression of wt KIF2A, but not KIF2A S100A, rescued both DKK1 and siLRP6 effects (Figure 4E,F and S3D). Furthermore, we partially knocked down KIF2A, which still resulted in congression defects (Figure 4G,H). In agreement with a role of mitotic Wnt signaling controlling KIF2A recruitment to the spindle, acute Wnt3a treatment in mitosis restored KIF2A levels at the spindle under low siKIF2A conditions, and rescued the congression defects (Figure 4G,H).

Finally, given the importance of Wnt signaling in stem cell proliferation and maintenance (Blauwkamp et al., 2012, Clevers et al., 2014), we investigated whether Wnt also promotes faithful execution of mitosis in pluripotent stem cells through the regulation of chromosome congression. Inhibition of basal Wnt signaling by DKK1 or the PORCN inhibitor LGK-974 led to a wide chromosome spread during metaphase in human induced pluripotent stem cells (hiPSCs) (Figure 5A,B). Importantly, ectopic expression of KIF2A rescued the effect of DKK1 in hiPSCs (Figure 5C,D). Taken together, these results indicate that Wnt signaling promotes chromosome congression and alignment by recruiting KIF2A to the spindle in somatic cells and pluripotent stem cells (Figure 5E).

**Figure 5.**
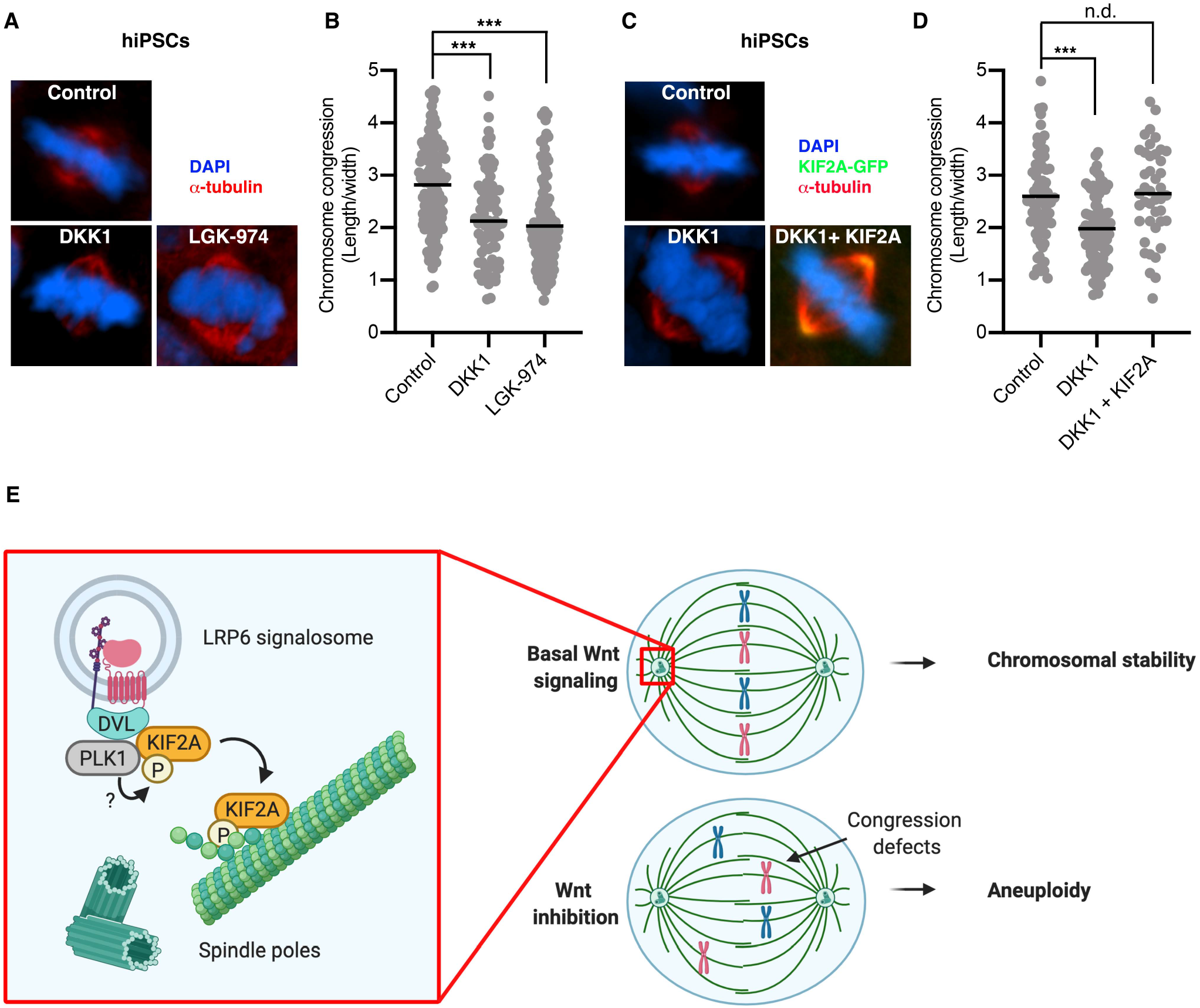
Wnt signaling promotes chromosome congression in hiPSCs. **A**,**C**, Representative immunofluorescence microscopy images of mitotic human induced pluripotent stem cells (hiPSCs) treated with the Wnt inhibitors DKK1 and LGK-974. In (**C**), cells were transfected with empty vector or KIF2A-GFP **B**,**D**, Quantification of the chromosome congression from (**A**,**C**). The length occupied by the chromosomes at the metaphase plate was divided by the width towards the spindle poles in n > 85 (**B**) and n > 42 (**C**) mitotic cells per condition. The median value is indicated. Representative images and analyses of n*≥* 3 independent experiments. **E**, Scheme of the suggested model for Wnt modulating chromosome congression in mitosis.

## DISCUSSION

The roles of Wnt signaling in G1/S progression are well established (Niehrs and Acebron, 2012), but its functions during cell division are poorly understood. The main conclusion of our study is the discovery that Wnt promotes chromosome congression and alignment during mitosis by controlling KIF2A localisation to the spindle poles. Our results support the following model: Wnt activity during G2/M promotes Dishevelled clustering in LRP6 signalosomes. Dishevelled recruits KIF2A through its motor and N-terminal domains, thereby facilitating its interaction with and activation by PLK1. Downstream of PLK1, phosphorylation of KIF2A at S100 by an unknown kinase promotes the interaction with Dishevelled and binding to the spindle. Finally, by monitoring KIF2A levels at the spindle poles, Wnt ensures proper chromosome alignment and prevents aneuploidy (Figure 5E).

Our results further highlight a critical role of canonical Wnt signaling in modulating the spindle dynamics during mitosis by different mechanisms. First, our previous research identified that Wnt/STOP signaling increases microtubule plus end dynamics, which leads to whole chromosome missegregation and aneuploidy (Stolz et al., 2015). Second, this study unravels a role of Wnt regulating KIF2A, a depolymerase functioning at the microtubule minus ends, i.e. at the spindle poles. Notably, both mechanisms converge to ensure proper chromosome alignment and segregation (Manning et al., 2007, Ertych et al., 2014, Ali et al., 2017, Trofimova et al., 2018). Interestingly, recent evidence on KIF18A shows that chromosome misalignment does not necessarily lead to aneuploidy (Fonseca et al., 2019). Hence, it would be important to further characterize the different processes controlled by KIF2A during mitosis and assess their relationship with CIN.

A standing question is whether the Wnt-mediated control of KIF2A plays a role in spindle scaling during development (Wilbur and Heald, 2013), especially considering that Wnt/STOP signaling is also required during the first embryonic cleavages (Huang et al., 2015). Furthermore, it is possible that these mechanisms may contribute to the Wnt conserved roles in spindle orientation and asymmetric cell division (Walston et al., 2004, Ciruna et al., 2006, Kikuchi et al., 2010, Sugioka et al., 2011, Habib et al., 2013).

Another important conclusion from our study is that Wnt signaling not only regulates proliferation and self-renewal (ten Berge et al., 2011, Blauwkamp et al., 2012), but also controls faithful execution of mitosis in pluripotent stem cells (Figure 5). Our research is consistent with -and suggests a mechanism for-previous evidence showing that Wnt secretion controls genome stability in mouse embryonic stem cells (Augustin et al., 2017). Interestingly, it has been shown recently that self-renewal of pluripotent cells is tightly linked to their genome integrity (Su et al., 2019). Hence, mitotic Wnt signaling may contribute to the known TCF/β-catenin functions in stem cell renewal by preventing chromosome instability.

Finally, given the critical roles of KIF2A in neurogenesis by modulating both cilium disassembly and neuronal wiring (Homma et al., 2003, Miyamoto et al., 2015, Ogawa and Hirokawa, 2015, Homma et al., 2018, Zhang et al., 2019), which are also key biological processes monitored by Wnt signaling (Hall et al., 2000, Sugioka et al., 2011, Weiner et al., 2020), it would be important to explore whether Wnt modulates KIF2A’s functions beyond mitosis.

## METHODS

### Cell culture

HeLa and HEK293T cells (ATCC) were cultured in DMEM medium (Gibco) supplemented with 10% FBS and 1% penicillin/streptomycin. RPE1-hTert cells (ATCC) were cultured in DMEM-F12 medium (Gibco) supplemented with 10% FBS, 1% penicillin/streptomycin and 4% Sodium Bicarbonate. All cells were grown at 37 °C with 5% CO_2_. HeLa cells, stably expressing H2B-mCherry and EGFP-α-tubulin, were a kind gift from Jan Ellenberg (Neumann et al., 2010). The human induced pluripotent stem cells were a gift from Kyung-Min Noh and were cultured in Essential E8 Medium (ThermoScientific), supplemented with Penicilin/Streptomycin (Chen et al., 2011). Revitacell Supplement (ThermoScientific) was added for the first 24h after plating). Medium was changed every day and cells were split every 4-5 days using versene solution. All the experiments were carried out with hiPSCs below passage 13.

Control and Wnt3a conditioned media were obtained from stably transfected L-cells. Dkk1 conditioned medium was produced by transiently transfecting HEK293T cells with pCS-FLAG-DKK1 plasmid using the Calcium-Phosphate method (15 µg plasmid per 10 cm dish). The medium was harvested 48 h after transfection.

To enrich cells in prometaphase, cells were first synchronized in G1/S for 24 h with 2 mM thymidine. After washing and releasing for 5 h in culture medium, cells were synchronized in mitosis using 2 µM dimethylenastron treatment for 4 h.

### siRNA transfection

Cells were transfected with 10-50 nM siRNAs (Sigma) using Lipofectamine RNAiMAX transfection reagent (Thermo Fisher Scientific). 48 h after transfection, cells were further analyzed. The following siRNAs were used: siRNA against Scramble (SIC001), KIF2A (SASI_Hs02_00319177, GAAGCUAUUCUUGAGCAAA), Mix of 1:1 LRP5/6 (SASI_Hs01_00086821, SASI_Hs01_00039493), Mix of 1:1:1 DVL1/2/3 (SASI_Hs01_00142403, SASI_Hs01_00142404, SASI_Hs01_0004203) and CTNNB1 (SASI_Hs01_00117960).

### DNA transfection and expression constructs

Cells were transfected with 250-800 ng DNA per well of a 12-well plate or 500 ng per well of a 6-well plate using X-tremeGENE 9 DNA transfection reagent (Roche), and harvested after 24 h.

pCS2+, pCS-FLAG-DKK1, FLAG-GSK3, FLAG-β-catenin, FLAG-Dvl1, FLAG-Dvl2, FLAG-Dvl3, FLAG-hCK1ε, hLRP6, mFzd8, hAxin1, Myc-Dvl2, TOPflash and Renilla plasmids were kindly provided by C. Niehrs. pEGFP-Kif2A (#52401), 3xFLAG-DVL2 (#24802), FLAG-Axin1 (#109370) and FLAG-β-TrCP (#10865) plasmids were obtained from Addgene. The KIF2A-phospho-mutant (KIF2A S100A) and the truncation mutants were generated by PCR with the following primers:

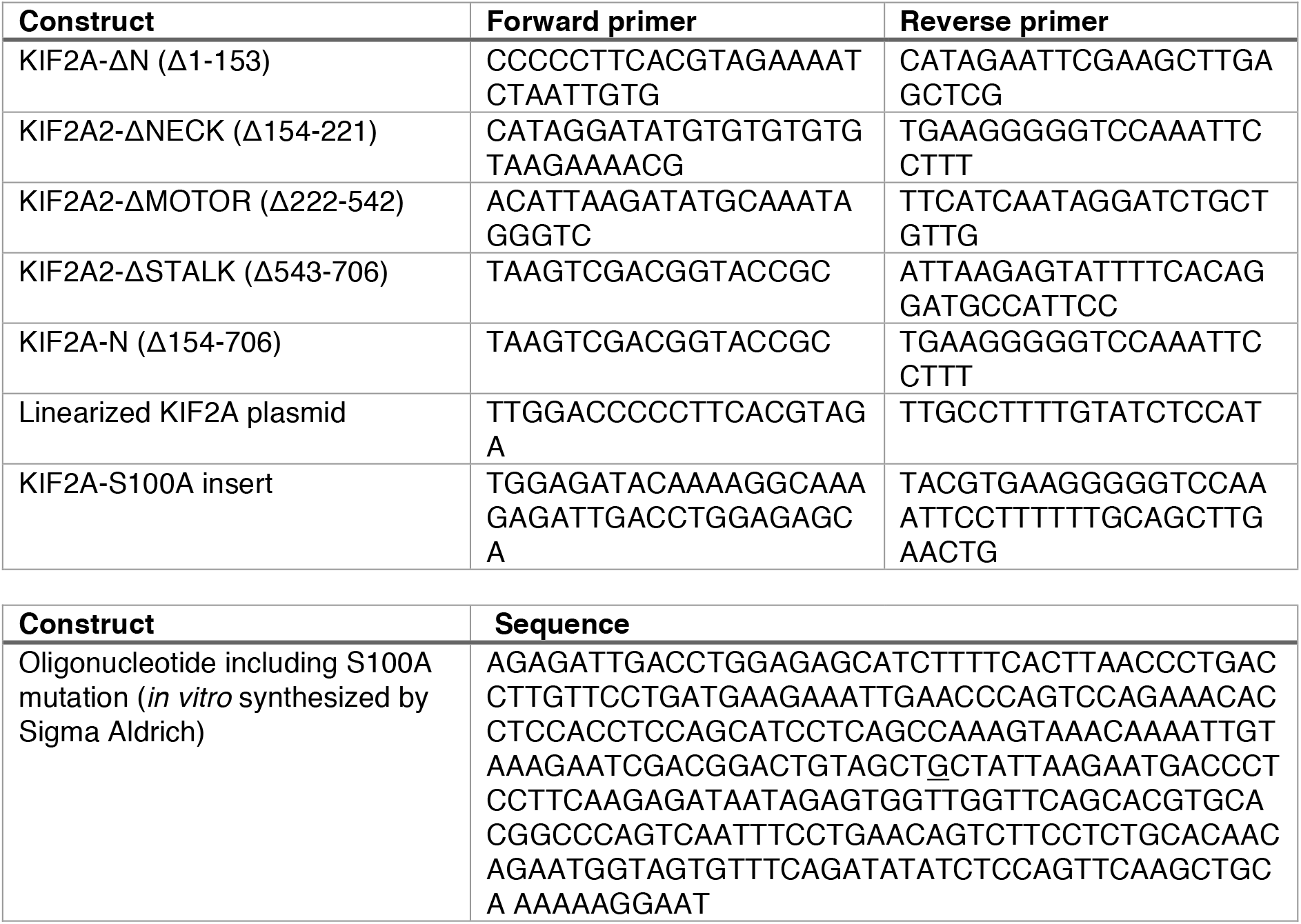

### MS sample preparation and analysis

RPE1 were cultured in either Medium or Heavy SILAC media (Cambridge Isotope Laboratories) as previously described (Ong et al., 2002). The experiment also included Light SILAC media labeled cells, which were not used for the downstream analysis. Cells were seeded in 15 cm dishes, synchronized in prometaphase, and treated with in-house produced control (Medium) or DKK1 (Heavy) SILAC media for 1.5 h before harvesting. Phosphoproteome analysis was performed as described previously (Borisova et al., 2017). Briefly, cells were lysed in modified RIPA buffer (50 mM Tris pH 7.5, 650 mM NaCl, 1 mM EDTA, 1% NP-40, 0.1% sodium deoxycholate) supplemented with protease inhibitors (Complete protease inhibitor cocktail tablets, Roche Diagnostics), 1 mM sodium orthovanadate, 5 mM β-glycerophosphate, and 5 mM sodium flouride (Sigma). After sonication, the lysates were cleared by centrifugation at 16,000 × *g* for 15 min and protein concentrations were estimated using QuickStart Bradford Protein assay (BioRad). After combining the same amount of proteins from each label, the proteins were precipitated in fourfold excess of ice-cold acetone and subsequently re-dissolved in denaturation buffer (6 M urea, 2 M thiourea in 10 mM HEPES pH 8.0). Cysteines were reduced with 1 mM dithiothreitol (DTT) and alkylated with 5.5 mM chloroacetamide. Proteins were digested with endoproteinase Lys-C (Wako Chemicals) and sequencing grade-modified trypsin (Sigma). Protease digestion was stopped by addition of trifluoroacetic acid to 0.5% and precipitates were removed by centrifugation. Peptides were purified using reversed-phase Sep-Pak C18 cartridges (Waters) and eluted in 50% acetonitrile. For the enrichment of phosphorylated peptides, 5 mg of peptides in binding buffer (50% acetonitrile, 6% trifluoroacetic acid in H_2_O) were incubated with 10 mg of TiO_2_ spheres (GL Sciences) for 1 h. The supernatant was used for a second enrichment step. The beads were washed twice in binding buffer, twice in wash buffer (0.5% TFA, 50% ACN) and subsequently peptides were eluted using elution buffer (10% NH_4_OH, 25% acetonitrile in H_2_O). The eluates were concentrated to remove NH_4_OH and peptides were fractionated in five fractions using micro-column-based strong-cation exchange chromatography and desalted on reversed-phase C18 StageTips (Weinert et al., 2013).

Peptide fractions were analyzed on a quadrupole Orbitrap mass spectrometer (Q Exactive Plus, Thermo Scientific) equipped with a UHPLC system (EASY-nLC 1000, Thermo Scientific) (Olsen et al., 2007). Peptide samples were loaded onto C18 reversed phase columns (15 cm length, 75 _m inner diameter, 1.9 _m bead size) and eluted with a linear gradient from 8 to 40% acetonitrile containing 0.1% formic acid in 2 h. The mass spectrometer was operated in data dependent mode, automatically switching between MS and MS2 acquisition. Survey full-scanMS spectra (*m/z* 300–1650) were acquired in the Orbitrap. The ten most intense ions were sequentially isolated and fragmented by higher energy C-trap dissociation (HCD) (Olsen et al., 2007). Peptides with unassigned charge states, as well as with charge states less than +2 were excluded from fragmentation. Fragment spectra were acquired in the Orbitrap mass analyzer.

Raw data files were analyzed using MaxQuant (development version 1.5.2.8) (Cox and Mann, 2008). Parent ion and MS2 spectra were searched against a database containing 92,578 human protein sequences obtained from the UniProtKB released in December 2016 using Andromeda search engine (Cox et al., 2011). Spectra were searched with a mass tolerance of 6 ppm in MS mode, 20 p.p.m. in HCD MS2 mode, strict trypsin specificity and allowing up to two miscleavages. Cysteine carbamidomethylation was searched as a fixed modification, whereas cysteine modification with n-ethylmaleimide, protein N-terminal acetylation, methionine oxidation, and phosphorylation of serine, threonine, and tyrosine were searched as variable modifications. Site localization probabilities were determined by MaxQuant using the PTM scoring algorithm as described previously (Cox and Mann, 2008). The dataset was filtered based on posterior error probability to arrive at a false discovery rate below 1% estimated using a target-decoy approach (Elias and Gygi, 2007). For downstream analysis, only phosphorylated peptides with a localization probability ≥ 0.75 were used. Fold-changes > 2 in both replicate experiments were considered significant.

### Immunofluorescence

RPE1 or HeLa cells were seeded on coverslips in 12-well plates. Cells were transfected with siRNA for 48 h, transfected with DNA for 24 h and treated with conditioned medium for 1.5 h, as indicated. Cells were fixed with 2% paraformaldehyde in PBS and stained, according to the experiment, for DAPI, α-tubulin, KIF2A, BUBR1 and CENPC.

The following antibodies were used: rabbit anti-α-tubulin (Sigma Aldrich, SAB4500087), mouse anti-KIF2A (Santa Cruz, sc-271471), mouse anti-BUBR1 (BD Bioscience, 612502), guinea pig anti-CENPC (MBL, PD03).

Coverslips were imaged with a Nikon Eclipse Ti using a 60x objective with oil immersion and the NIS Elements software. Data was analyzed using ImageJ 2.0.0. To access KIF2A levels at the spindle, a threshold for the KIF2A signal was set to only detect KIF2A at the poles.

### Real-time PCR

Cells were synchronized as described above, treated with control, Dkk1 or Wnt3a conditioned medium for 1.5 h and harvested. Total cellular RNA was extracted by using the RNeasy Mini Kit (QIAGEN) and cDNA was synthesized from 1 μg of total RNA using the Bioline SensiFAST cDNA Synthesis Kit. Quantitative PCR was performed with the SensiFAST SYBR Hi-ROX Kit (Bioline) using a StepOnePlus 96-well plate reader (Applied Biosystems). Gene expression was normalized to GAPDH.

The following primer pairs were used for qPCR

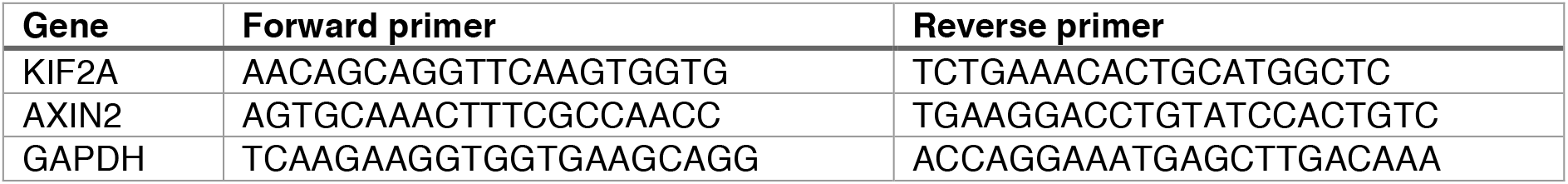

### Western blotting

For Western blotting, cells were lysed in full lysis buffer (50 mM Tris-HCl, pH 7.5, 150 mM NaCl, 1% NP-40, 0.05% SDS, 1 mM β-mercaptoethanol, 2 mM EDTA, 1x protease phosphatase inhibitor cocktail (Thermo Fisher)) or cytoplasmic lysis buffer (PBS supplemented with 0.05% saponin, 10 mM β-mercaptoethanol, 2 mM EDTA, 1x protease phosphatase inhibitor cocktail). The cleared lysates were mixed with 4x NuPAGE loading buffer, resolved on 8% NuPAGE gels and transferred to nitrocellulose membranes. For western blot experiments the following antibodies were used: mouse anti-α-tubulin (Sigma Aldrich, T9026), mouse anti-β-catenin (BD Bioscience, 610153), mouse anti-KIF2A (Santa Cruz, sc-271471), rabbit anti-LRP6 (Cell Signaling, 2560S), rabbit anti-DVL (Merck Millipore, ABD122), chicken anti-GFP (Abcam, ab13970), mouse anti-FLAG M2 (Sigma Aldrich, F1804).

### Proximity ligation assay

HeLa cells were seeded on coverslips in 12-well plates. Cells were transfected with siRNA for 48 h, transfected with DNA for 24 h and treated with conditioned medium for 1.5 h as indicated. Cells were fixed with 2% paraformaldehyde in PBS and proximity ligation assays (PLA) were performed with the Duolink In Situ Red Starter Kit Mouse/Rabbit (Sigma Aldrich) according to the manufacturer’s protocol.

For the overexpression PLAs, EGFP-KIF2A and a FLAG-tagged component were overexpressed and the following antibodies were used: rabbit anti-GFP (Abcam, ab290), mouse anti-FLAG M2 (Sigma Aldrich, F1804). For the endogenous PLAs mouse anti-KIF2A (Santa Cruz, sc-271471), rabbit anti-DVL (Merck Millipore, ABD122), and rabbit anti-PLK1 (Abcam, ab14209) were used.

Coverslips were imaged with a Nikon Eclipse Ti using a 60x objective with oil immersion and the NIS Elements software. Data was analyzed using ImageJ 2.0.0. The PLA signal was quantified by the number of PLA complexes per cell (Figure 2C,H,J) or the total fluorescence signal per cell (Figure 2F). Values are displayed relative to the control.

### Wnt reporter assays (TOPflash)

HEK293T or HeLa cells were seeded into 96-well plates. HEK293T cells were transfected with 50 ng DNA per well, including 5 ng TOPflash luciferase and 3 ng Renilla luciferase. HeLa cells were transfected with 100 ng of DNA containing 30 ng TOPflash luciferase, 20 ng Renilla luciferase, and co-transfected, where indicated, with 25 ng Myc-DVL2 or 25 ng Myc-DVL2, 22 ng LRP6, 2 ng mouse Fzd8 and 1.25 ng AXIN1. The transfection was performed using X-tremeGENE 9 DNA transfection reagent (Roche) according to manufacturer instructions. After 24 h, cells were treated overnight as indicated, and lysed with 1x Passive Lysis Buffer (Promega) on a shaker for 15 min at 4 °C. Lysates were analyzed using a Dual-Luciferase Reporter Assay System (Promega) with a Tecan Microplate Reader (Tecan Infinite M1000). TOPflash signals were normalized to the Renilla reporter and Wnt activity was calculated relative to the control condition.

### Karyotype analysis

RPE1 cells were transfected every 5 days with 10 nM siRNA against KIF2A, LRP6 and Scrambled (Control) over 30 days and karyotype analyses was performed as described (Stolz et al., 2015). In brief, cells were treated with 2 μM dimethylenastron for 4 h to enrich for mitotic cells. Cells were pelleted, washed with PBS and incubated in hypotonic medium (40% DMEM-F12, 60% H_2_O) at RT for 15 min. Cells were fixed in Carnoy’s solution (methanol:acetic acid = 3:1). Chromosomes were spread onto glass slides and stained with Giemsa solution. Chromosome number variability was determined by chromosome counting.

### Live cell imaging

HeLa cells stably expressing H2B-mCherry and EGFP-α-tubulin were seeded in a 10-well CELLview slide (Greiner Bio-One). Before imaging, medium was replaced by DMEM medium with HEPES and without phenol red (Gibco) supplemented with 10% FBS, 1% penicillin/streptomycin an 1% sodium pyruvate. Live cell imaging was performed using an automated Nikon Eclipse Ti2 inverted microscope equipped with a 20x water immersion objective (NA 0.95) and a sNEO CMOS camera (Andor). Multipoint acquisition was controlled by NIS-Elements 5.1 software. Image stacks were recorded every 10 min for up to 14 h in a chamber (STXG-WSKM, Tokai Hit) at 37 °C and 5% CO_2_. Images were analyzed using ImageJ 2.0.0 software.

### FACS

RPE1 cells were synchronized in mitosis and treated for 1.5 h with control SILAC Medium or Dkk1 SILAC Heavy media. Cells were harvested and fixed for FACS analysis measuring the DNA content (propidium iodide) and mitotic phospho-epitopes (MPM-2) as described (Ertych et al., 2014).

### Determination of microtubule plus-end assembly rates

Microtubule plus end growth rates were determined by tracking EB3-GFP in living cells as described previously (Ertych et al., 2014). Briefly, cells were transfected 48 h prior to the measurement with siRNA as indicated and pEGFP-EB3 (kindly provided by L. Wordeman, University of Washington, USA), seeded onto glass bottom dishes (Ibidi) and treated with 2 μM Dimethylenastron for 2 h before measurement. Live-imaging was performed using a Deltavision ELITE microscope (GE Healthcare) with a 60x objective. Images were recorded every 2 seconds while cells were incubated at 37 °C and 5% CO_2_. Images were deconvolved using SoftWorx 5.0/6.0 software and average assembly rates were calculated from 200 individual microtubules (20 individual microtubules per cell. n=10 cells).

### Image data processing

Raw images were imported to Fiji (ImageJ, v2.0) prior to their export to Photoshop 2020 for panel arrangement. Linear changes in contrast or brightness were equally applied to all controls and across the entire images. The models and schemes were created with BioRender.com.

### Statistical analyses

Data are shown as mean with standard error of the mean (SEM or SD), as indicated in the figure legends. Where indicated, Student’s t-tests (two groups) or one-way ANOVA analyses with Tukey correction (three or more groups) were calculated using Prism v8. Significance is indicated as: *P < 0.05, **P < 0.01, ***P < 0.001, or n.s.: not significant.

## ACKNOWLEDGMENTS

We thank C. Niehrs, K.M. Noh, J. Ellenberg, and T.W. Holstein for sharing reagents. We thank the Nikon Imaging Center and the FACS core facility at the University of Heidelberg for access to microscopes and cytometers, as well as for technical help. This work was supported by the Deutsche Forschungsgemeinschaft (DFG) through the SFB1324. A.d.J.S. holds a Humboldt Research Fellowship for Postdoctoral Researchers.

## AUTHOR CONTRIBUTIONS

H.B. and S.P.A. conceived the project. A.B., A.G.A. Y.C., M.I.H., A.C., A.d.J.S, and S.P.A. conceived, performed and analysed experiments. M.O. and P. B. performed and analysed the SILAC experiment. U.E. assisted with the live cell imaging experiments. S.P.A. supervised all aspects of the project. S.P.A. and A.B. wrote the manuscript with input from all authors.

## FIGURE LEGENDS

**Figure S1.**
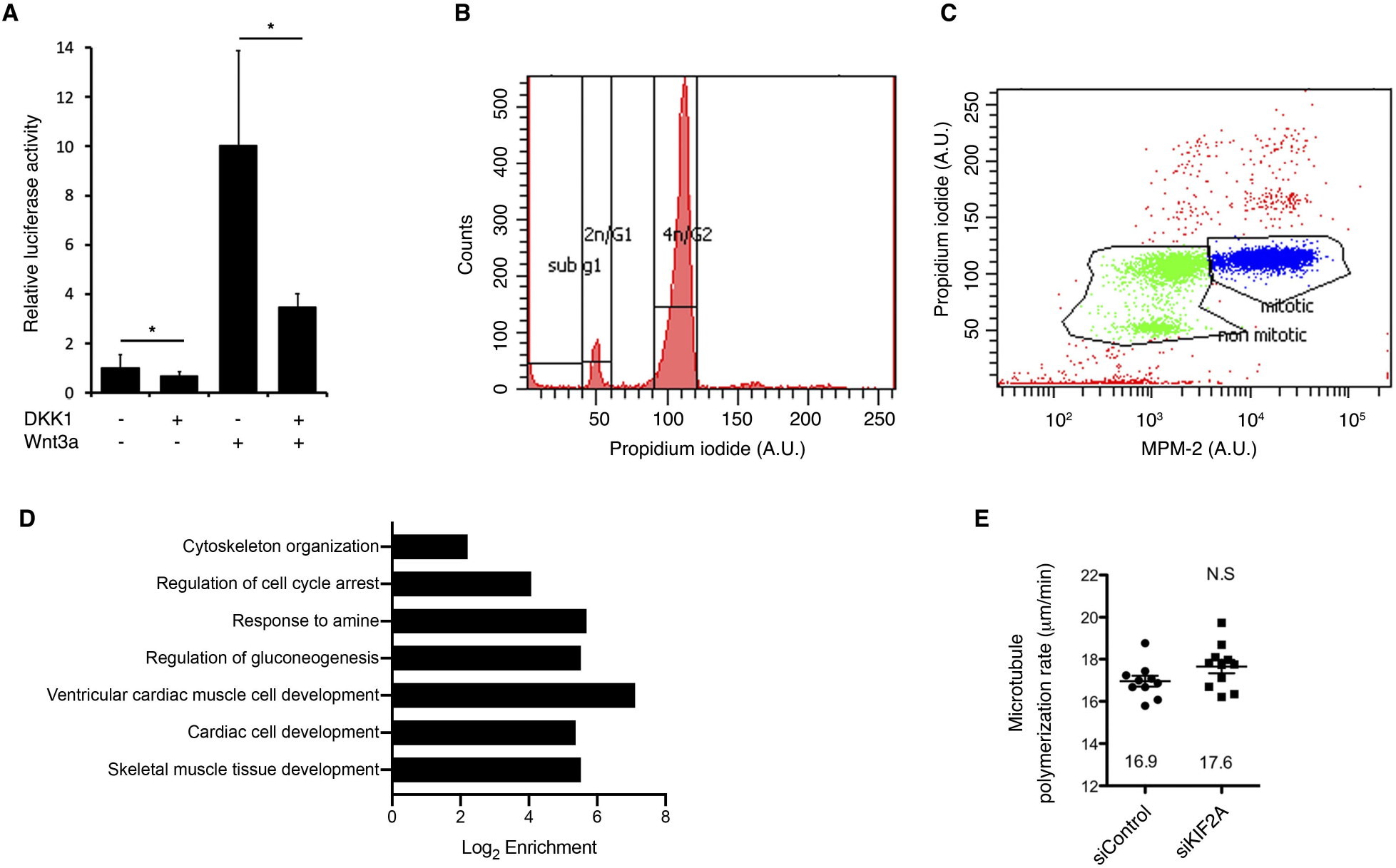
Controls for the mitotic analyses of Wnt signaling and KIF2A. **A**, Wnt reporter assays showing the activity of DKK1 and Wnt3a conditioned media used in these studies. Data is displayed as mean ± SD of 3 biological replicates. **B**,**C**, Cell cycle profiles of cells that were synchronized as described in Figure 1A. Mitotic cells (PI = 4n; MPM-2 positive) are shown in blue and represent *∼*66% of the population. **D**, Gene ontology enrichment (Log_2_) of DKK1 target proteins (Fold change > 2 in each experiment) from Figure 1A,B. **E**, Measurements of microtubule plus end assembly rates in EB3-GFP stably transfected RPE1 cells during mitosis after siRNA-mediated knockdown of KIF2A (20 microtubules/cells, n = 10 cells).

**Figure S2.**
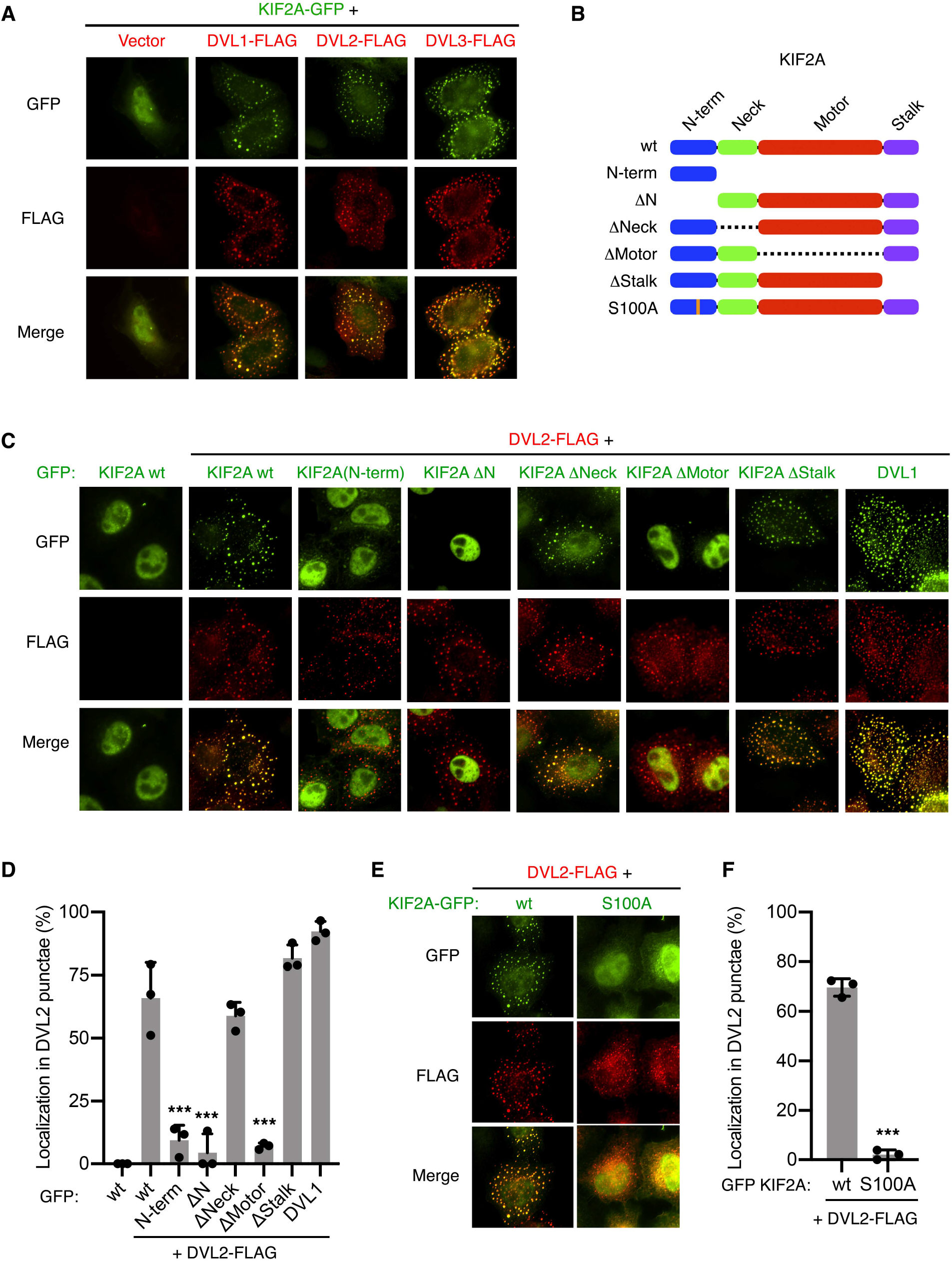
KIF2A is recruited by Dishevelled through its N-terminal and motor domains. **A**,**C**,**E**, Representative immunofluorescence microscopy images from n ≥ 3 independent experiments showing HeLa cells transfected with the indicated FLAG- or GFP-tagged constructs. **B**, Scheme showing the KIF2A constructs used in (**C-F**). **D**,**F**, Bar plots showing the quantification of cells with strong co-localization of GFP-tagged constructs and DVL2-FLAG from (**C**,**E**). Data is displayed as mean + S.D. percentage of cells from n = 3 independent experiments with n ≥ 20 cells/condition in each independent experiment.

**Figure S3.**
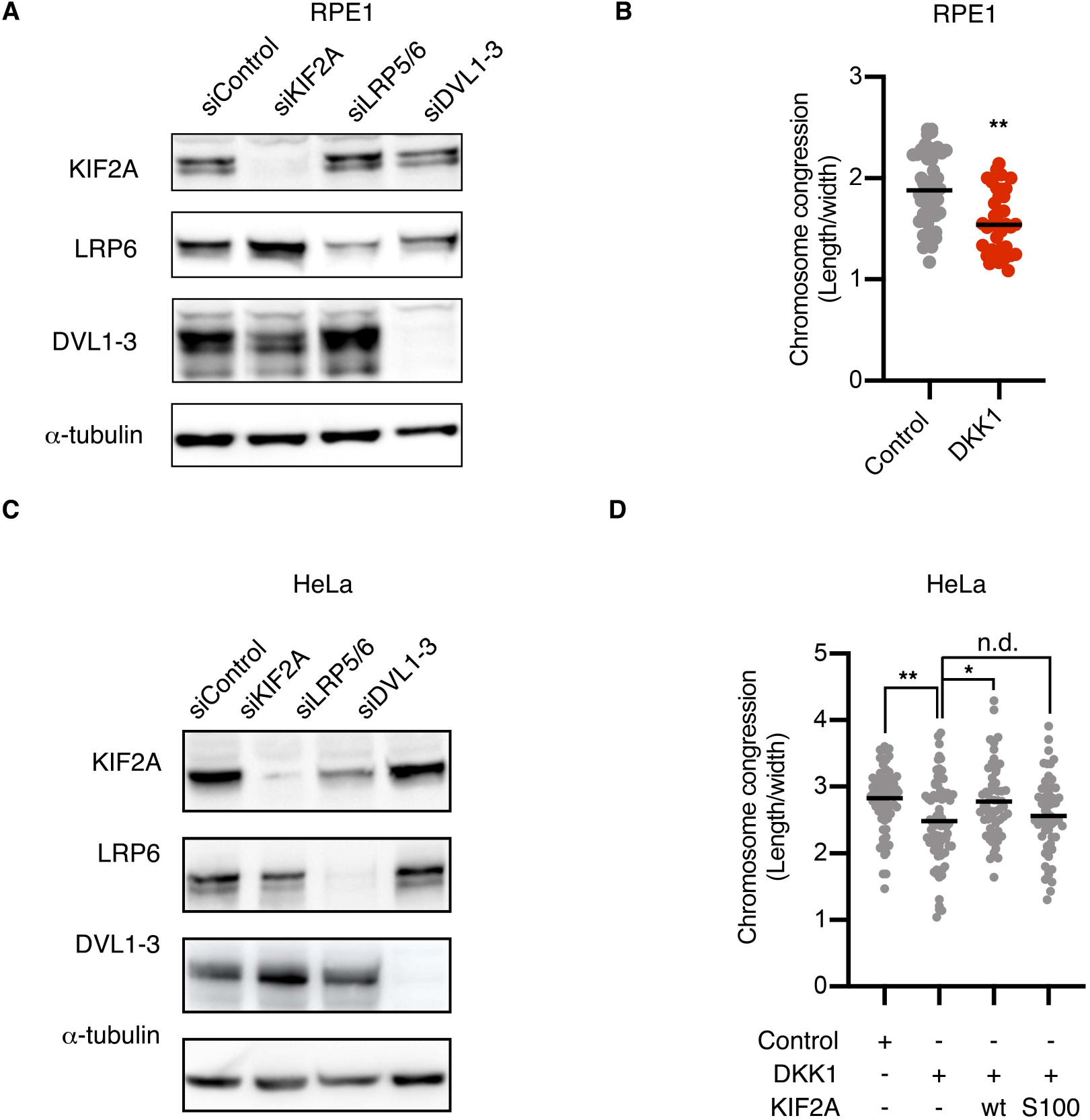
Wnt promotes chromosome congression in RPE and HeLa cells. **A**,**C** Western blots showing the knockdown efficiency of the siRNAs in RPE1 and HeLa cells shown in Figures 3,4, respectively. **B**,**D**, Quantification of chromosome congression in metaphase of RPE1 cells (**B**, median of n > 30 cells per condition) and HeLa cells (**D**, median of n > 52 cells per condition) treated with control or DKK1 conditioned media for 1.5h. In (**D**) cells were transfected with KIF2a wt, KIF2A S100A, or empty vector. Representative analyses of n ≥ 3 experiments.

